# Typhoid toxin hijacks Wnt5a to potentiate TGFβ-mediated senescence and *Salmonella* infections

**DOI:** 10.1101/2022.10.05.510870

**Authors:** Mohamed ElGhazaly, Mark O Collins, Angela EM Ibler, Daniel Humphreys

## Abstract

Damage to our genome causes acute senescence in mammalian cells, which undergo growth arrest and release a secretome that elicits cell cycle arrest in bystander cells through the senescence-associated secretory phenotype (SASP). Thus, acute senescence is a powerful tumour suppressor. *Salmonella enterica* hijacks senescence through its typhoid toxin, which usurps unidentified factors in the stress secretome of senescent cells to mediate intracellular infections. Here, transcriptomics of toxin-induced senescent cells (txSCs) and proteomics of their secretome identified secreted ligands that activate the TGFβ pathway through SMAD transcription factors. The ligand Wnt5a established a self-amplifying positive feedback loop driving TGFβ signalling, which enforced autocrine senescence in txSCs and paracrine senescence in naive bystander cells by activation of DDRs. Wnt5a and GDF15 increased host cell susceptibility to infection. The study reveals how an innate defence against cancer is co-opted by a bacterial pathogen to cause widespread damage and mediate infections.

## Introduction

Damage to our genomes can generate double-strand DNA breaks (DSBs) resulting in hereditable gene mutations and cells with oncogenic potential. To counteract pathology, the DNA damage response (DDR) is activated through kinases ATM (ataxia-telangiectasia mutated), which responds to DSBs, and ATR (ATM and rad3-related) that senses single-strand DNA breaks (SSBs), e.g. generated during DNA replication stress whereupon ATR phosphorylates RPA(1). Uniting the functions of ATM and ATR is phosphorylation of their effector, histone H2AX at S139 (γH2AX)(2, 3), which recruits and retains DDR proteins at sites of damage to coordinate repair and cell fate decisions including survival, apoptosis and senescence(1, 4–6).

Cellular senescence is a hallmark of ageing that restricts the growth of cells with genomic stress through stable cell cycle arrest(7, 8). Senescent cells display persistent γH2AX and 53BP1 DDRs, elevated expression of cyclin-dependent kinase inhibitors, down-regulation of the nuclear envelope component lamin B1 and an increase in lysosomal content marked by senescence-associated-β-Galactosidase (SA-β-Gal)(7, 8). Senescent cells release a signature pro-inflammatory secretome that establishes the senescence-associated secretory phenotype (SASP)(8–10). SASP factors modulate the microenvironment to enforce senescence responses in the senescent cells in an autocrine fashion while establishing paracrine senescence in naive bystander cells(9, 11, 12). The factors also recruit immune cells to eliminate damaged cells with pathological potential(13–15). Consequently, acute senescence is a powerful tumour suppressor(7, 8). However, persistent DDRs cause chronic SASP and the resulting inflammation promotes age-related pathologies such as cancer. For example, persistent DNA damage caused by the toxin colibactin of *Escherichia coli* induced SASP that promoted colon tumour growth(16). While the DDR is best known for its ability to prevent proliferation of cancerous cells(6), the DDR continues to be revealed as an important innate defence against infections by intracellular bacteria(17, 18). Thus, bacterial pathogens have evolved sophisticated ways to manipulate host DDRs and execute their virulence strategies(7, 19). This is exemplified by genotoxic serovars of *Salmonella enterica*, particularly the human pathogen *S*.Typhi, which causes 14 million cases of acute typhoid fever per year and asymptomatic, often life-long, chronic infections in individuals who have increased risk of gallbladder carcinoma(20).

To cause typhoid and chronic disease, *S*.Typhi establishes intracellular intestinal infections and evades innate immune responses before disseminating to systemic sites(20). Intracellular *S*.Typhi reside within the *Salmonella*-containing vacuole (SCV) from where it expresses the typhoid toxin comprising PltB-PltA-CdtB subunits that is exocytosed into the extracellular milieu(21). Once deployed, the PltB subunit binds to sialylated glycans on host surface receptors facilitating toxin endocytosis(22). Reduction of disulphide bonds linking PltA-CdtB liberates the toxigenic DNase1-like subunit CdtB, which translocates to the nucleus where it activates DDRs through nuclease activity(21–24). The toxin is also encoded by related *Salmonella* serovars including *S*.Paratyphi and *S*.Javiana that cause disease in humans and food-chain animals worldwide(25, 26). Toxin-induced DDRs are implicated in *Salmonella* evasion of intestinal inflammatory responses(26, 27), typhoid fever(22, 28), bacteraemia(28), dissemination(26, 27), and chronic infection(27). The mechanisms by which toxin-induced DDRs mediate *Salmonella* infections are unclear.

Typhoid toxin was recently shown to induce acute cellular senescence in cultured fibroblasts(23), as well as the intestinal mucosa and T lymphocytes of infected mice(24, 29). Senescence was caused by two distinct DDRs: DSBs marked by ATM-dependent canonical γH2AX foci and SSBs signified by ATR-dependent pan-γH2AX localisation throughout the nucleus, which was enriched at the nuclear periphery marking a response <u>in</u>duced by a <u>g</u>enotoxin (RING)(23). *Salmonella* hijacked senescence to promote intracellular infections. This was dependent upon an unidentified toxin-induced secretome (secretome-TxWT) released from senescent cells, which transmitted senescence to bystander cells indicative of toxin-associated SASP (txSASP). Unlike viruses, pathogenic bacteria do not depend on host cell proliferation to replicate so how senescence might counteract infection is unclear. Instead, senescence may enable subversion of the host secretome that can remodel bystander cells in the infection niche(7).

The human host secretome plays a critical part in determining the outcome of bacterial infections(30–33). However, host secretome responses to a bacterial toxin have not yet been resolved. Here, we sought to determine the identity of the toxin-induced host secretome from senescent cells underlying txSASP to further address the significance of senescence in bacterial infections. The study reveals secreted ligands that converge on the transforming growth factor beta (TGF-β) pathway, which generated a self-amplifying autocrine loop enforcing DDRs in intoxicated cells while causing paracrine senescence in non-stimulated bystander cells. The secreted txSASP factors increased host cell susceptibility to *Salmonella* infection.

## Results

### The DNA damage response secretome from cells undergoing acute senescence

Unidentified factors in the secretome of host cells intoxicated for 48h were shown to elicit txSASP - paracrine senescence and enhanced infection in bystander cells(23). To identify the factors underlying txSASP, we first generated toxin-induced senescent cells (TxSC) as depicted in **Fig S1A**: HT1080 fibroblasts were intoxicated for 2h with 20ng/ml of wild-type purified recombinant typhoid toxin (TxWT), which resulted in stalled synthesis of EdU-labelled DNA due to cell-cycle arrest (**Fig S1B**) and entry into a senescence-like phenotype at 96h marked by up-regulation of SA-β-Gal (**Fig S1C, S1D**). This contrasted with untreated control cells (unt) or cells treated with toxin lacking DNase activity due to a H160Q mutation in CdtB (TxHQ) that progressed through the cell-cycle and lacked SA-β-Gal (**Fig S1C, S1D**). We found that TxSCs continued to adopt signatures of senescence including persistent γH2AX responses (**Fig S1E-S1H**). As illustrated in Fig 1A, TxSC secretomes were harvested at 24h intervals (secretome-TxWT) before assaying SA-β-Gal activity in naive bystander HT1080 fibroblasts (**Fig 1B**). Relative to the secretomes from untreated cells (secretome-Unt), all batches of secretome-TxWT induced paracrine senescence, which increased from 60% of cells with batch 0-24h to a maximum of 75% of cells with batches 24-48h, 48-72h and 72-96h (**Fig 1B**). Combining all batches of secretome-TxWT had no additive effect. This indicates that mediators of paracrine senescence in secretome-TxWT released from TxSCs undergoing acute senescence responses reached a maximum threshold by 48h.

**Figure 1.**
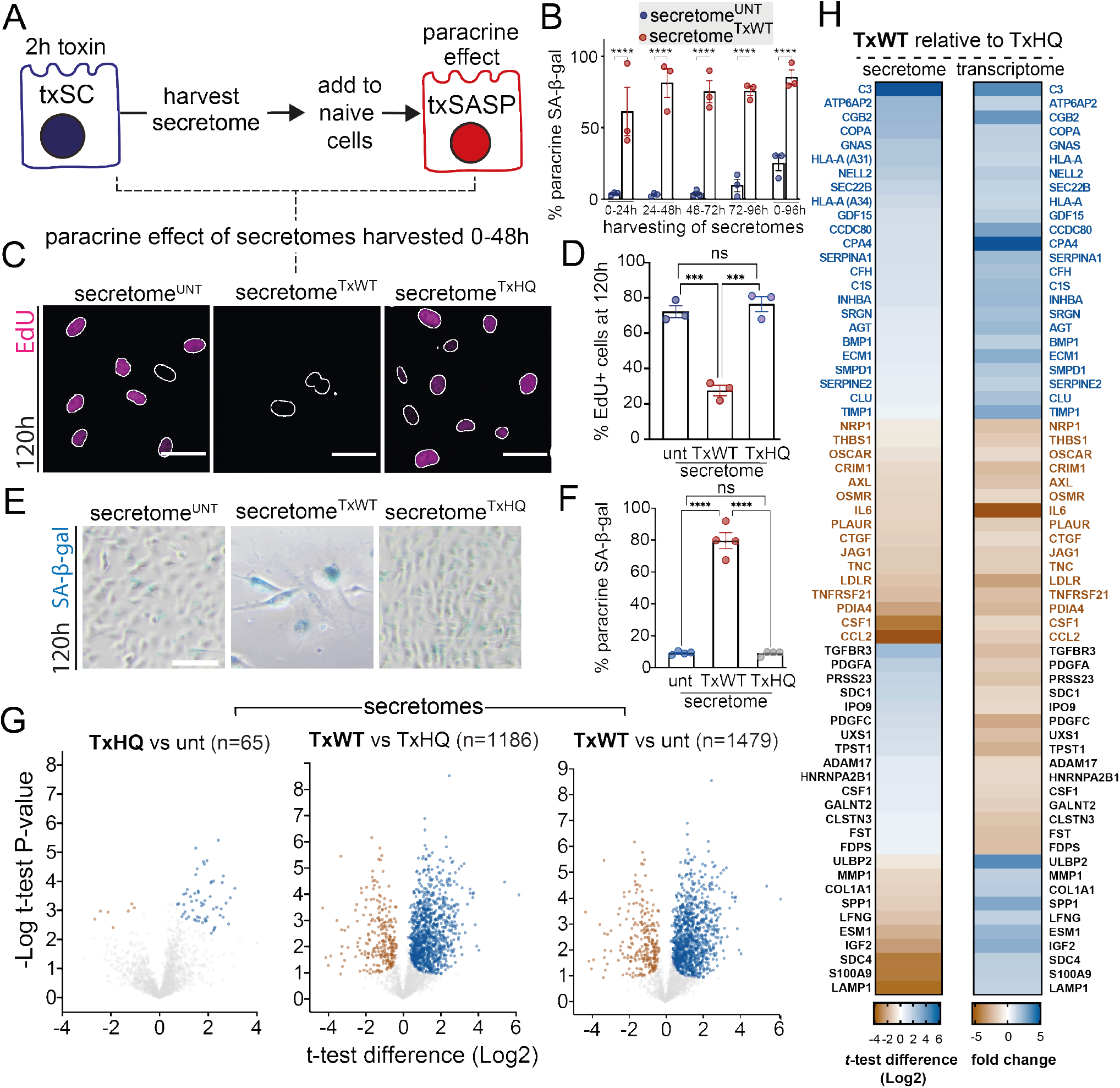
The typhoid toxin-induced host secretome underlying txSASP. **(A)** Experimental schematic depicting txSASP mediated via secretomes harvested from TxSCs. **(B)** Percentage of SA-β-gal-positive naive HT1080 cells at 96h following treatment with conditioned media containing secretomes harvested at indicated intervals from untreated (secretome^UNT^) or TxWT-treated (secretome^TxWT^). Circles represent technical replicates (1 biological replicate). **(C)** Representative images of naive cells undergoing replication of EdU-labelled DNA (magenta) 120h after incubation with secretomes from untreated (secretome^UNT^), TxWT-treated (secretome^TxWT^) or TxHQ-treated (secretome^TxHQ^) cells harvested between 0-48h. Outlines of DAPI-stained nuclei indicated. **(D)** Quantification of (C). Circles represent biological replicates. **(E)** SA-β-gal in cells from experiment described in (C) with quantification in **(F)** where circles represent biological replicates. **(G)** Volcano plots indicating significantly upregulated (blue dots) and down-regulated (brown) proteins in secretomes harvested as described in (C). Perseus two-sample t-test (s0=0.01 and FDR=0.05) from 4 biological repeats. **(H)** Heat map analysing gene expression of factors in TxWT-induced secretome relative to TxHQ-treated cells at 48h determined by GeneChip microarray transcriptome analysis (p<0.05). Blue text indicates upregulated, brown downregulated, and black text indicates opposing fold changes between analyses. For microscopy images, scale bars 50 μm in (C) and (E). Asterisks indicate significance (*p<0.05, **p<0.01, ***p<0.001, ****p<0.0001, ns=no significance) relative to untreated unless indicated otherwise calculated by (B) Two-way ANOVA, (D) and (F) One-way ANOVA, with Tukey’s multiple comparison test. **Fig 1** connected to **Fig S1 - S3**.

To identify the proteome of secretome-TxWT, we pooled batches of secretome-TxWT harvested between 0-48h (workflow depicted in **Fig S2A**) before validating paracrine senescence in naive cells by assaying induction of cell-cycle arrest (**Fig 1C, 1D**), SA-β-Gal (**Fig 1E, 1F, S2B–2C**) and DDRs marked by γH2AX and 53BP1 (**Fig S2D–2H**). Next, quantitative LC-MS/MS enabled unbiased identification of proteins in secretome-Unt, secretome-TxHQ and secretome-TxWT (**Fig 1G**). There was very little difference between negative controls secretome-Unt and secretome-TxHQ (n=65 significant proteins), which contrasted with secretome-TxWT that was significantly divergent from secretome-Unt (n=1479) and secretome-TxHQ (n=1186). The majority of significant proteins were up-regulated in secretome-TxWT (**Fig 1G;** 954 proteins out of 1186 indicated with blue circles in TxWT vs TxHQ), which was also observed in the secretome of cells undergoing irradiation- and oncogene-induced senescence (OIS)(34). No typhoid toxin (i.e. CdtB-PltA-PltB) was identified in the proteomic analysis (**Fig S2I**), though we observed the non-toxigenic subunit PltB, which was presumably bound to recycled receptors shed from the cell surface. **Fig S2J** reveals secretome-TxWT from TxSCs comprising 149 proteins, which were defined by their presence within the human secretome(35) and their differential regulation relative to negative controls (P< 0.05, n=149 significant proteins).

### Toxin-induced host secretome implicates TGFβ signalling in cellular senescence

To further define secretome-TxWT, the transcriptome of TxSCs was determined by GeneChip microarray transcriptome analysis of cells at 48h following 2h treatment with TxWT or control TxHQ (**Fig 1H**). Of the 149 proteins comprising secretome-TxWT (**Fig S2J**), 24 proteins were identified as significant by both proteomics and transcriptomics (**Fig 1H**). Secretome-TxWT included known SASP proteins such as effectors of innate immunity (e.g. complement C3, C1r and C1s) that were up-regulated in both analyses (indicated by **blue** labelling), as were SERPIN-A1, ECM1 and TIMP1 that were identified in the SASP Atlas database(34). Of the proteins that were down-regulated in both proteomic and transcriptomic analyses (**brown** labelling), the most notable was interleukin-6 (IL6), which is implicated in SASP(7, 8, 10). TGFβ-ligands INHBA (henceforth Activin A), GDF15 and BMP1 that activate ALK receptors were found up-regulated in both analyses (**Fig 1H**). Interestingly, inhibitors of Activin A-driven TGFβ signalling, TGFBR3 and follistatin (FST), were found up-regulated by proteomics but down-regulated by transcriptomics (**black** labelling), perhaps suggesting that the receptor TGFBR3, for example, is shed from TxSCs rather than exhibiting increased expression. Indeed, the sheddase ADAM17 was also up-regulated, which is implicated in liberating cell surface-bound SASP proteins during OIS(8).

We next determined whether txSASP is transmitted through up-regulated or down-regulated proteins in secretome-TxWT. When we replenished the down-regulated proteins in secretome-TxWT by supplementing with secretome-Unt or secretome-TxHQ, elevated SA-β-Gal was still observed indicating paracrine senescence is mediated via up-regulated proteins in secretome-TxWT (**Fig S3A, S3B**). This reinforced our interest in TGFβ-family ligands up-regulated in secretome-TxWT (e.g. Activin A, GDF15, BMP1 in **Fig 2H**). Indeed, PANTHER gene-function analysis identified TGFβ signalling as the most significantly up-regulated pathway in secretome-TxWT relative to secretome-TxHQ (**Fig S3C**).

**Figure 2.**
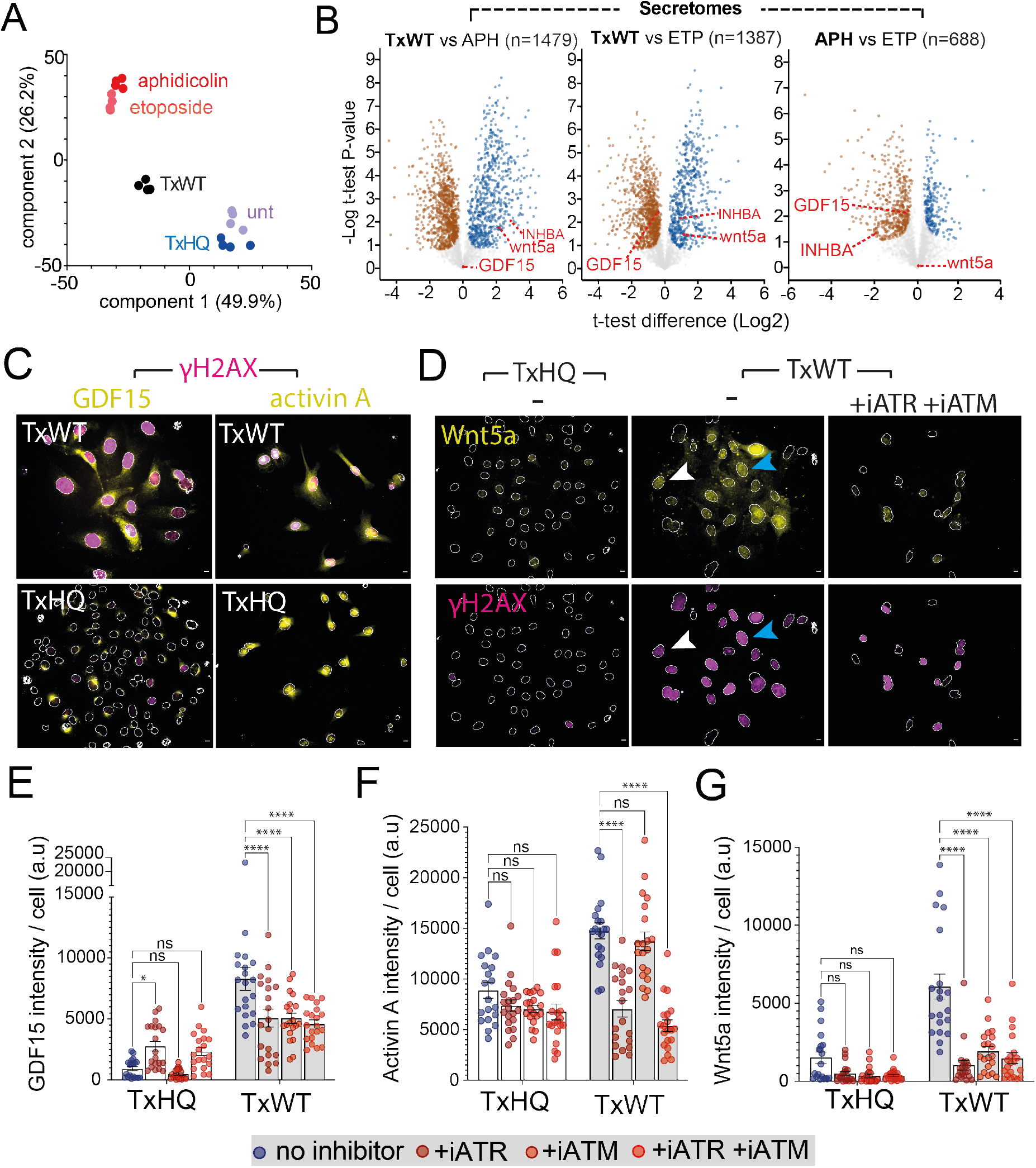
Regulation of txSASP by ATM and ATR. (**A**) Principal component analysis of secretomes from untreated, TxWT-, TxHQ-, aphidicolin- or etoposide-treated cells harvested between 0-48h post chase. (**B**) Volcano plots indicating significantly upregulated (blue dots) and downregulated (brown) proteins in secretomes harvested as described in (A). Perseus two-sample t-test (s0=0.01 and FDR=0.05) from 4 biological repeats. Red labels indicate Wnt5a, INHBA (Activin A) and GDF15. (**C**) Representative images of γH2AX (magenta), GDF15 and Activin A (yellow) in TxWT- or TxHQ-treated cells at 48h. Outlines of DAPI-stained nuclei indicated. Scale bars are 10 μm. (**D**) Same experiment as (C) assaying Wnt5a expression −/+ inhibitors of ATM and ATR. (**E**-**G**) Quantification of GDF15, Activin A, and Wnt5a expression in TxWT- or TxHQ-treated cells at 48h −/+ inhibitors of ATM and ATR. Circles represent technical replicates from 2 biological repeats. Asterisks indicate significance (*p<0.05, **p<0.01, ***p<0.001, ****p<0.0001, ns=no significance) relative to no inhibitor controls calculated by Two-way ANOVA with Dunnett’s multiple comparisons test. Error bars SEM. **Fig 2** connected to **Fig S4**.

### Typhoid toxin induces expression of txSASP factors Wnt5a, Activin A and GDF15

TGFβ signalling pathway represents a major axis in oncogene-induced transmissible senescence (8, 12, 14, 36). Thus, our data indicate a conserved role for TGFβ ligands in senescence triggered by divergent DNA damage inducers that include typhoid toxin. To investigate this further, we treated cells with TxWT, the SSB-inducer aphidicolin (APH) or the DSB-inducer etoposide (ETP)(1) before examining senescence responses. Like TxWT, we found that APH and ETP both induced senescence-like responses marked by γH2AX and SA-β-Gal at 96h (**Fig S3D**), and paracrine senescence in ∼40% of naive cells demonstrating SASP (**Fig S3E**). LC-MS/MS and principal component analysis (PCA) of the secretomes revealed 3 distinct clusters of senescence-inducers APH and ETP (red), negative controls secretomes (unt and TxHQ in blue), which were distinct from TxWT (black) that formed an independent cluster (**Fig 2A**). We found secretome-TxWT was significantly divergent from secretome-APH (1479 proteins) and secretome-ETP (1387 proteins) while secretome-APH and secretome-ETP had more overlap with only 688 differentially regulated proteins (**Fig 2B**).

To identify conserved mediators of paracrine senescence, the secretomes of all senescence inducers (TxWT, APH and ETP) were filtered against the untreated negative control (**Fig S3F**). The TGFβ family member GDF15 was immediately striking (**Fig S3F**), which was also identified in the TxWT-induced secretome (**Fig 1H**). GDF15 is differentially expressed during ageing, negatively regulates IL6 expression, and was identified as a SASP component promoting colon cancer (37–39).

To identify constituents specifically enriched in txSASP, secretome-TxWT was filtered against the secretomes from both negative (unt, TxHQ) and positive controls (APH, ETP) (**Fig S3G**). We found regulators of TGFβ signalling INHBA (i.e. Activin A), TGFBR3, FST, the transmembrane protein vasorin (VASN) and metalloprotease ADAM17 (TGFβ signalling summarised in cartoon **Fig S4**). However, though significantly up-regulated in secretome-TxWT across all conditions, Activin A (labelled red) was also found up-regulated in secretome-ETP relative to secretome-Unt and -APH (**Fig 2B**) suggesting a conserved role for Activin A in senescence. In the search for unique constituents in secretome-TxWT, we identified Wnt5a (labelled red in **Fig 2B**). Thus, Wnt5a is an enriched component of txSASP that also contains common SASP factors Activin A and GDF15. Taken together, the data implicates TGFβ signalling and Wnt5a in a stress secretome from cells undergoing senescence-like responses due to typhoid toxin.

### TxSASP is orchestrated by ATM and ATR

To understand how txSASP is established, we examined expression of Activin A, GDF15, and Wnt5a at 96h in cells treated for 2h with TxWT or the nuclease-deficient derivative TxHQ (**Fig 2C, 2D**). Nuclease activity of TxWT induced expression of Activin A, GDF15, and Wnt5a, which was coincident with γH2AX (**Fig 2C-2G**). Previously, typhoid toxin was shown to cause ATM-dependent γH2AX-foci that labelled DSBs in the G1 phase of the cell cycle and ATR-dependent RPA foci, which labelled SSBs marked by pan-γH2AX in G2(23). Thus, ATM and ATR respond to distinct DDRs induced by typhoid toxin. Indeed, the distinct DDRs were observed in **Fig S1E** where intoxicated cells displaying γH2AX foci (white arrows) lacked phosphorylated RPA (pRPA) while nuclei with pan-γH2AX contained pRPA (blue arrows). To dissect the role of ATM and ATR in txSASP, cells were intoxicated in the presence of small molecule inhibitors of ATM (iATM) and ATR (iATR). We first focussed on Wnt5a, which was specifically enriched in secretome-TxWT (**Fig 2B, S3G**). We found that Wnt5a was coincident with TxWT-treated cells containing γH2AX foci (white arrow) and pan-γH2AX (blue arrow) (**Fig 2D**). Consistent with this, Wnt5a expression was significantly reduced in the presence of iATR or iATM, and when both inhibitors were applied in combination (**Fig 2D, 2G**). The same trend was observed for GDF15 (**Fig 2E**). In contrast, TxWT-induced Activin A expression was reliant on ATR but not ATM (**Fig 2F**), which indicated a prominent role for ATR in driving expression of TGFβ ligands and Wnt5a.

### Toxin activation of TGFβ signalling elicits γH2AX responses and senescence

Elevated TGFβ signalling has been previously observed in mice infected with *Salmonella* Typhimurium(40) but no role in the host-pathogen interaction has been ascribed. TGFβ and Activins activate Type 1 ALK tyrosine kinase receptors (ALK1-7) that phosphorylate SMAD2/3 transcription factors while BMPs tend to activate SMAD1/5/8, both of which, drive gene transcription in diverse cellular pathways including senescence(36)(summarised in cartoon **Fig S4**). Robust phosphorylation of SMAD2 (pSMAD2) at 48h was evident with TxWT in a nuclease-dependent manner as TxHQ was equivalent to untreated controls (**Fig 3A, 3B**). TxWT-dependent pSMAD2 signalling was also observed in HIEC6 intestinal epithelial cells, IMR90 and NIH3T3 fibroblasts (**Fig S5A, S5B, S5C**). TGFβ ligands were observed secreted in response to TxWT, APH and ETP (**Fig 2B**). Consistent with a conserved role for TGFβ ligands in paracrine senescence (**Fig 2B**), APH and ETP triggered pSMAD2 production, which was also observed with the positive control TGFβ3 (**Fig 3A, 3B**). The same trend was observed in SMAD1/5, which was activated by TxWT, APH, ETP but not TxHQ (**Fig S5D, S5E**). We found that infection of host cells by wild-type (WT) toxigenic *Salmonella* Javiana induced wide-spread γH2AX, which was coincident with activation of SMAD2 at 48h and 96h (**Fig 3C, 3D**). SMAD2 activation was not observed with *ΔcdtB S*.Javiana confirming a requirement for toxin-induced DDRs. The data show activation of SMAD transcription factors downstream of TGFβ-family ligands identified in secretome-TxWT.

**Figure 3.**
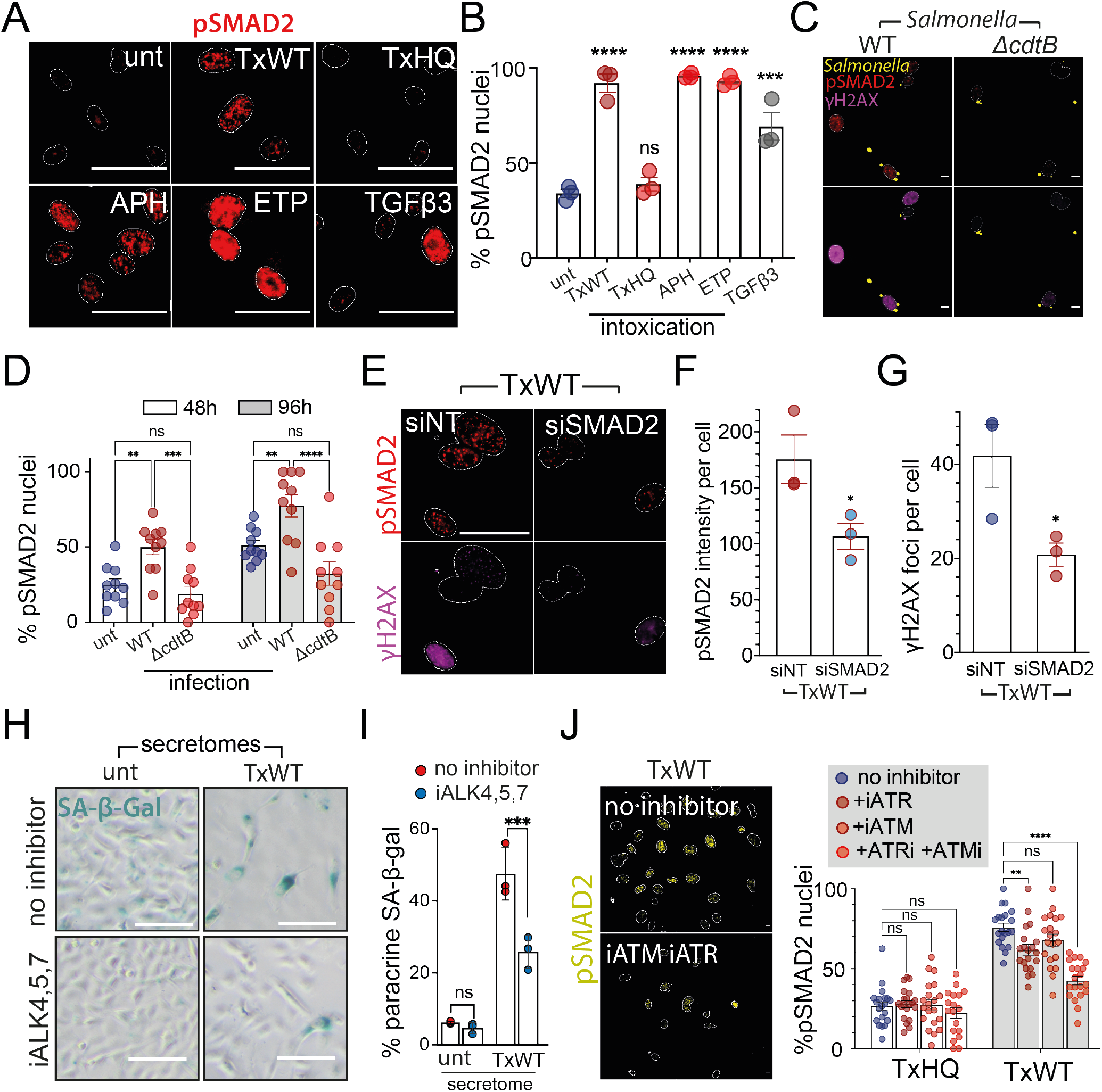
Toxin manipulation of TGFβ-mediated host DDRs and senescence. (**A**) Representative images of pSMAD2 at 48h in cells with indicated treatments. (**B**) Quantification of (A) relative to the average pSMAD2 integrated intensity of untreated cells (circles indicate 3 biological replicates). (**C**) Representative images of pSMAD2 (red) and γH2AX (magenta) cells at 96h following 30 min infection with wild-type (WT) or *ΔcdtB S*.Javiana (yellow). (**D**) Quantification of the experiment in (C) at 48h and 96h. Circles represent technical replicates (1 biological replicate). (**E**) Representative images of pSMAD2 (red) and γH2AX (magenta) in TxWT-treated cells 48h after non-targeting (siNT) or SMAD2 (siSMAD2) siRNA transfection. Quantification of the experiment in (E) assaying (**F**) pSMAD2 integrated intensity per nucleus and (**G**) number of γH2AX foci per nucleus. Circles indicate 3 biological replicates. (**H**) Representative images of SA-β-gal in cells at 120h treated with secretome^UNT^ or secretome^TxWT^ from Fig 1C with or without the inhibitor of ALK4,5,7. (**I**) Quantification of the experiment in (H). (**J**) Representative images of pSMAD2 (left) in TxWT-treated cells −/+ ATM and ATR inhibitors at 48h with quantification (right) where circles indicate technical replicates (2 biological replicates). For images in (A), (C), (E), and (J), outlines of DAPI-stained nuclei are indicated. Scale bars are 50 μm (A, E) or 10 μm (C, J). Asterisks indicate significance (*p<0.05, **p<0.01, ***p<0.001, ****p<0.0001, ns=no significance) calculated by (B) One way ANOVA with Tukey’s multiple comparisons test, (D, I) Two-way ANOVA with Sidak, (J) Dunnett’s multiple comparisons test, and (F-G) unpaired t-test. Error bars SEM. **Fig 3** connected to **Fig S5**.

Previous studies showed that replicative-, oncogene- and drug-induced SASP activates DDRs in bystander cells to perpetuate senescence through the TGFβ pathway(36). This suggests that TGFβ-dependent signalling mediated through secretome-TxWT activates DDRs thereby contributing to txSASP. Thus, we inhibited pSMAD2 signalling by siRNA transfection (siSMAD2) before treatment with TxWT (**Fig 3E, 3F, 3G**). Activation of pSMAD2 was still observed in TxWT-treated cells transfected with non-targeting siRNA (siNT) but this was significantly reduced with siSMAD2 (**Fig 3E, 3F**). We found that the number of γH2AX foci per cell were reduced by 50% in SMAD2-depleted cells indicating that TGFβ signalling enforces senescence in TxSCs through DDRs **(Fig 3E, 3G)**. Moreover, a small molecule inhibitor of ALK4,5,7 significantly reduced paracrine senescence in cells treated with secretome-TxWT (**Fig 3H, 3I**). Taken together, the data indicate that typhoid toxin hijacks TGFβ signalling to enforce autocrine senescence in TxSCs while also mediating paracrine senescence in naive bystander cells impacted by secretome-TxWT thereby establishing txSASP.

### Wnt5a crosstalk with TGFβ pathway establishes txSASP

Inhibition of ATM and ATR in TxWT-treated cells undergoing cellular senescence impaired expression of TGFβ ligands and Wnt5a (**Fig 2D-2G**), which also inhibited activation of SMAD (**Fig 3J**). We thus sought to determine which component of secretome-TxWT contributes to SMAD2 activation. We initially investigated the role of Wnt5a in txSASP by depleting Wnt5a expression during 72h siRNA transfection (siWnt5a) before examining DDRs and paracrine senescence (**Fig 4**). In cells transfected with a non-targeting control (siNT), TxWT induced Wnt5a expression, which was significantly diminished by siWnt5a transfection (**Fig 4A, 4B**). To our surprise, Wnt5a depletion also reduced γH2AX (**Fig 4C**), which phenocopied the effect of SMAD2-depletion on γH2AX responses in **Fig 3E, 3G**.

**Figure 4.**
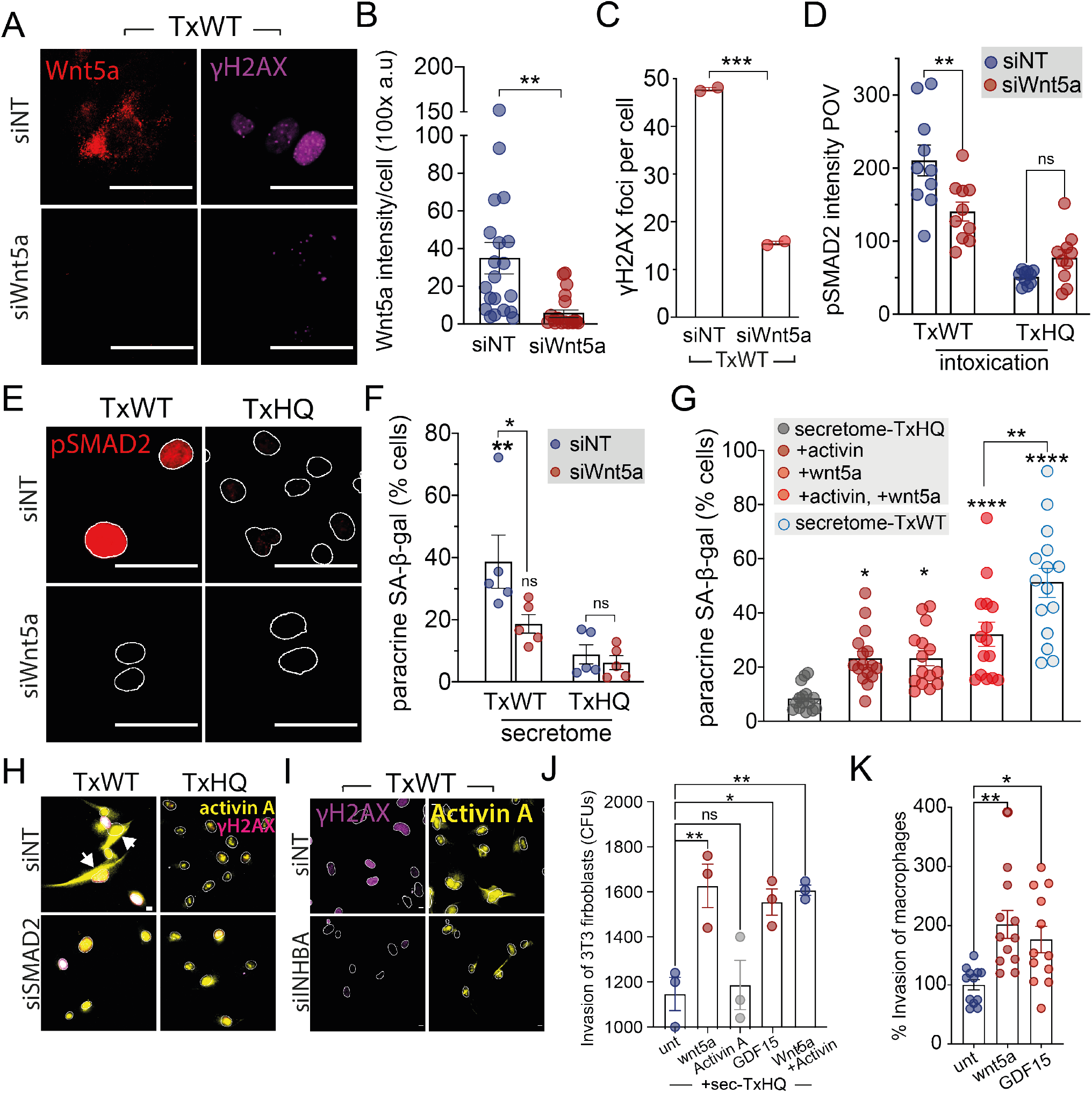
Wnt5a and TGFβ interplay underlying txSASP. (**A**) Representative images of Wnt5a (red) and γH2AX (magenta) in TxWT-treated cells 48h after non-targeting (siNT) or Wnt5a (siWnt5a) siRNA transfection. Quantification of (A) assaying: (**B**) Wnt5a knockdown with circles indicating technical replicates, and (**C**) number of γH2AX foci per nucleus with circles indicating 2 biological replicates. (**D**) Quantification of pSMAD2 intensity per field of view (POV) from the experiment in (**E**) that shows representative images of pSMAD2 in TxWT- or TxHQ-treated cells at 48h following transfection with siNT or siWnt5a. (**F**) Quantification of SA-β-gal at 120h in naive cells treated with secretome^TxWT^ or secretome^TxHQ^ harvested from the experiment in (E). Circles indicate 5 biological replicates. (**G**) Quantification of SA-β-gal at 48h in naive cells treated with secretome^TxHQ^ alone or supplemented with combinations of purified Activin A and Wnt5a. Secretome^TxWT^ used as positive control. Circles indicate technical replicates (3 biological replicates). Representative images of Activin A (yellow) and γH2AX (magenta) in TxWT-treated cells at 48h following transfection with siNT and siSMAD2 in (**H**), or siNT and siINHBA in (**I**). (**J**) *Salmonella* invasion into non-phagocytic cells. Colony forming units (CFUs) cultured from NIH3T3 fibroblasts infected for 30 mins in the presence of secretome^TxHQ^ alone (unt) or supplemented with purified Wnt5a, Activin A and GDF15. Circles represent technical replicates (1 biological replicate). (**K**) *Salmonella* uptake into THP1 macrophages in the presence of purified Wnt5a or GDF15 Circles represent technical replicates (2 biological replicates). For images in (A), (E), (H), and (I), outlines of DAPI-stained nuclei are indicated. Scale bars are 50 μm (A, E) or 10 μm (H, I). Asterisks indicate significance (*p<0.05, **p<0.01, ***p<0.001, ****p<0.0001, ns=no significance) calculated by (B-C) Unpaired t-test, (D) two-way ANOVA with Sidak multiple comparisons test, and (F-G, J) One-way ANOVA with Tukey’s and (K) Dunnett’s multiple comparison test. Error bars SEM. **Fig 4** connected to **Fig S5**.

The findings suggest Wnt5a might enforce senescence through SMAD2-dependent DDRs. Indeed, in control cells treated with TxHQ, Wnt5a depletion had no effect on activation of TGFβ signalling marked by pSMAD2 (**Fig 4E**). In contrast, Wnt5a depletion impaired TGFβ signalling in cells undergoing TxWT-induced senescence indicating a synergy between non-canonical Wnt5a and the TGFβ pathway. Consistent with TGFβ-dependent senescence (**Fig 3H, 3I**), Wnt5a depletion resulted in impaired paracrine senescence via secretome-TxWT (**Fig 4F**). To examine Wnt5a-TGFβ synergy during paracrine senescence, we incubated cells with control secretome-TxHQ with or without purified Wnt5a or Activin A (**Fig 4G**). Secretome-TxHQ alone caused no paracrine senescence, which was significantly increased with either Wnt5a or Activin A that was further enhanced when supplied in combination revealing an additive effect. Interestingly, secretome-TxWT caused more widespread paracrine senescence than the combination of Wnt5a or Activin A supporting the view that txSASP is established through multiple factors acting together, i.e. in addition to Wnt5a and Activin A. As Wnt5a activated SMAD2 (**Fig 4E**), we examined Wnt5a-TGFβ synergy in TxSCs by studying expression of Activin A (**Fig 4H**). We found that SMAD2-depletion abrogated Activin A expression (**Fig 4H**: white arrows). SMAD2 activation in TxSCs had no effect on the expression of Wnt5a and GDF15 demonstrating that only Activin A is downstream of pSMAD2 (**Fig S5F, S5G, S5H**). This suggests TxWT-mediated activation of SMAD2 elicits DDRs through Activin A. Indeed, TxWT-induced γH2AX γH2AX was impaired i γH2AX n Activin A-depleted cells γH2AX (**Fig 4I:** siINHBA). This enables us to propose that Wnt5a activates expression of Activin A via SMAD2 in a positive feedback loop, which acts in synergy to drive autocrine DDRs in TxSCs and establish paracrine senescence in by-stander cells.

### Wnt5a and GDF15 enhance host cell invasion by *Salmonella*

*Salmonella* invades host cells by promoting cytoskeletn remodelling that drives pathogen uptake(41). We previously showed that txSASP promotes *Salmonella* invasion into fibroblasts and macrophages through unidentified factors in secretome-TxWT(23). We thus examined the significance of Wnt5a-TGFβ-mediated txSASP γH2AX. First, we examined invasion into fibroblasts in the presence of negative control secretome-TxHQ alone (unt) or secretome-TxHQ supplemented with recombinant txSASP factors Wnt5a, Activin A or GDF15 (**Fig 4J**). We found that Wnt5a but not Activin A enhanced *Salmonella* invasion. Wnt5a and Activin A synergised to cause paracrine senescence (**Fig 4G**) but this was not observed when *Salmonella* invasion was examined in the presence of both Wnt5a and Activin A (**Fig 4J**). This suggests invasion is independent of SMAD-mediated signal transduction. For example, TGFβ can activate regulators of the actin cytoskeleton independently of SMADs(42) while Wnt5a promotes Rac1-mediated actin assembly(43). This view was supported by observations in infected cells treated with siRNAs targeting ALK2,4,5, or 7 receptors where only ALK4 showed a significant, yet modest, role in *Salmonella* uptake (**Fig S5I**). Like Wnt5a, GDF15 also promoted *Salmonella* invasion into fibroblasts (**Fig 4G**). Moreover, we found that Wnt5a and GDF15 also promoted *Salmonella* uptake into macrophages (**Fig 4K**).

Taken together, the data elucidates the secretome of host cells undergoing acute senescence responses to typhoid toxin, which establishes txSASP by amplifying DDRs in TxSCs causing paracrine senescence and enhanced infection through TGFβ ligands and crosstalk from Wnt5a.

## Discussion

SASP is considered a double-edged sword capable of exerting the beneficial effects of senescence (e.g. embryonic development, wound healing and tumour suppression) or its deleterious effects (e.g. cancer progression, ageing)(7, 8). In situations where the proliferation of cancer or virus-infected cells can drive disease, SASP factors that can reinforce and spread senescence in autocrine and paracrine mechanisms provides a powerful defence(7, 8, 13-15). However, bacterial pathogens are not reliant on mammalian DNA replication machinery for proliferation and SASP consequently provides a possible hijack target during infection. In this study, we resolved the identity of a host secretome released in response to genotoxic stress from typhoid toxin, which resulted in paracrine senescence and enhanced infections through Wnt5a potentiation of the TGFβ pathway - a mediator of senescence(36)(**Fig 5 - model**).

**Figure 5.**
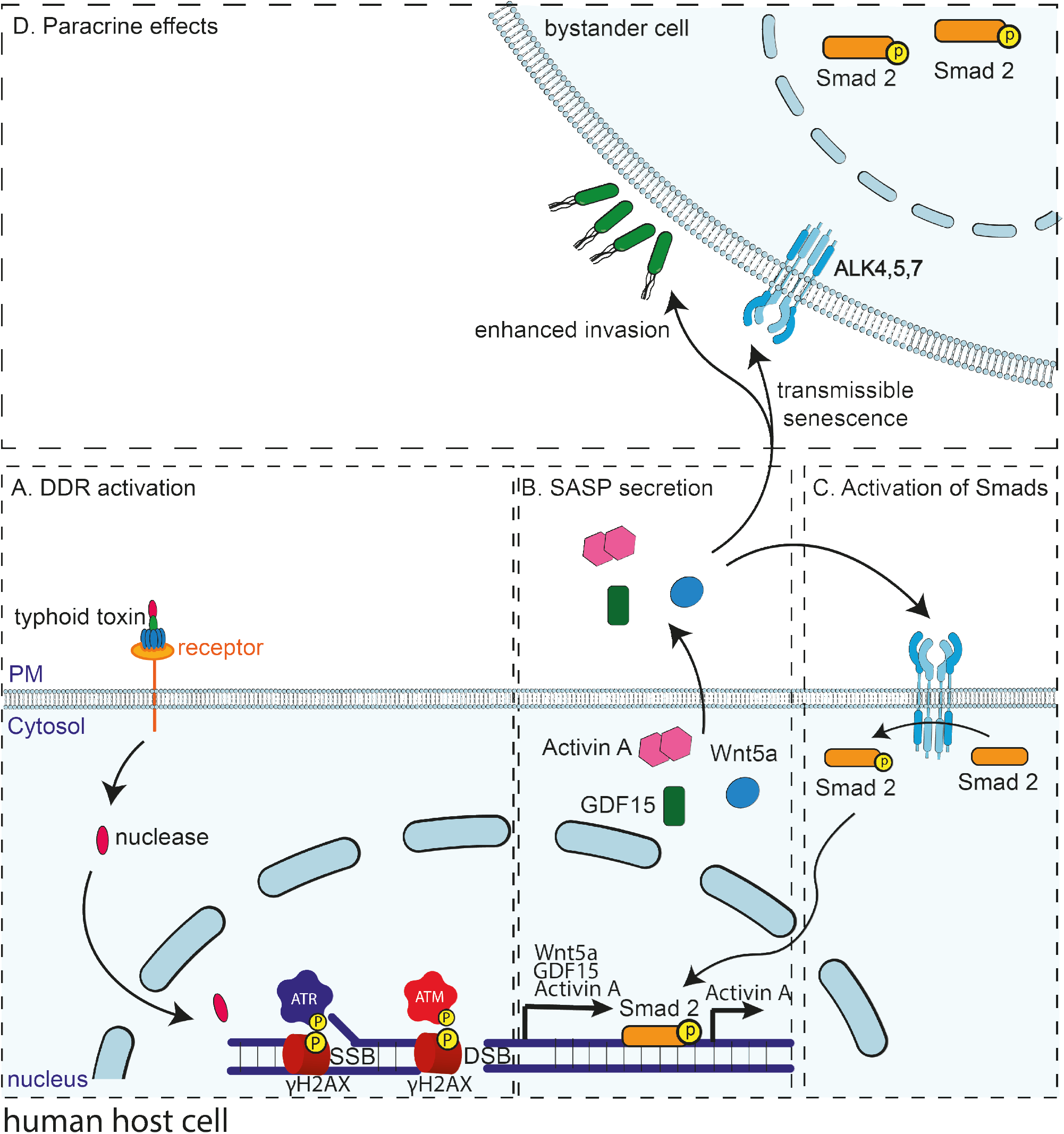
Proposed model. (**A**) The typhoid toxin activates DDRs through nuclease activity causing single- (SSB) and double-strand DNA breaks (DSB) resulting in activation of ATR and ATM leading to γH2AX. (**B**) ATR/ATM drive expression of Wnt5a, GDF15 and Activin A, which drive (**C**) autocrine SMAD signalling in TxWT-treated cells undergoing TxSC, and (**D**) paracrine SMAD signalling via ALK4,5,7 receptors and infection in naive bystander cells.

A role for TGFβ in senescence induced by bacterial pathogens has not been previously described. Consistent with this role, SMAD2/3 activation by TGFβ ligands induce cell cycle arrest by repressing proliferation factors while increasing expression of CDK inhibitors(44). TGFβ signalling also induces reactive oxygen species and DNA damage(44). Indeed, we found that Wnt5a, SMAD2 and Activin A amplified γH2AX DDRs in TxSCs, which has been observed with Activin A during OIS(12, 45, 46). Besides autocrine effects on TxSCs, we also found that Activin A was sufficient to promote paracrine senescence in naive cells, an activity which was enhanced by Wnt5a. This synergy was apparent in TxSCs, which was initiated by ATM/ATR-dependent expression of Wnt5a, Activin A and GDF15 (**Fig 5A, 5B**). Wnt5a activated SMAD2, which augmented expression of Activin A indicating a positive feedback loop that reinforces senescence in an autocrine manner via DDRs while amplifying TGFβ-mediated paracrine senescence in bystander cells (**Fig 5C, 5D**). In support of this mechanism, Wnt5a has been shown to potentiate the activity of TGFβ1 during regeneration of damaged colonic crypts in mice(47). This study consequently expands the synergistic role of Wnt5a and TGFβ to pathogen-induced senescence responses.

The human host secretome plays a critical part in determining the outcome of bacterial infections. For example, the *Salmonella* virulence effector protein SIfA causes exocytosis of antimicrobial cargo destined for the SCV, which promotes intracellular survival of *Salmonella* (30). On the other hand, host secretion can counteract infectious disease: bacterial toxins inactivate Rho GTPases, which is sensed by the host inflammasome as a decline in Rho activity resulting in secretion of IL-1β and IL-18 that recruit immune cells to limit infection(32). Here, we identify Wnt5a and GDF15 as secreted txSASP factors that enhance *Salmonella* infection. Bacterial hijacking of GDF15 has not been previously observed. Wnt5a has been shown to promote phagocytic uptake of *Escherichia coli* into macrophages(48), as well as *Streptococcus pneumoniae* and *Pseudomonas aeruginosa* where Wnt5a was also implicated in bacterial clearance(43). Thus, Wnt5a can act in host defence. However, *Salmonella* survives within macrophages and is thus able to hijack Wnt5a for invasion through a toxin-induced host secretome.

We found that typhoid toxin induces acute senescence responses from as early as 48h. SASP undergoes dynamic changes through its temporal regulation(10): ‘initial SASP’ comprises DDR-associated secretions from cells entering a senescence-like state and is followed by ‘early SASP’, which develops over several days and is associated with self-amplifying autocrine loops that reinforce senescence. ‘Late SASP’ reaches maturity after 4-10 days. In this study, we focussed on initial SASP where txSASP factors were known to mediate decisive processes involved in establishing infection(23). Even so, we revealed txSASP factors in secretome-TxWT that amplified senescence responses, which is indicative of early SASP(10).

## Conclusion

*Salmonella* exploits chronic senescence in ageing organisms, *indirectly*, during colonisation of systemic sites(49), chronic *Salmonella* carriage(50), and pathogen entry into senescent fibroblasts from elderly volunteers(51). This is consistent with the view that senescence increases susceptibility to infection in ageing organisms(7). Our findings indicate that *Salmonella* employs typhoid toxin to *directly* accelerate ageing-like responses through acute senescence, which may be significant *in vivo*. During infection of mice, *Salmonella* has been shown to induce senescence in a toxin-dependent manner resulting in suppression of inflammatory responses(24, 29). This is consistent with up-regulation of TGFβ ligands in txSASP and down-regulation of interleukin-6, which is an inflammatory mediator of SASP(8). The same observations were made during OIS where SASP enriched in TGFβ1/3 suppressed immunosurveillance of cancer cells(8, 14). Thus, our study advances the senescence field through an emerging theme investigating acute senescence responses to pathogenic bacteria, which has revealed a novel *Salmonella* hijack mechanism dependent on Wnt5a potentiation of the TGFβ pathway. The findings are of significance to those studying interplay between innate immunity and senescence, as well as those investigating how diverse bacterial pathogens exploit host DDRs to cause disease.

## Acknowledgements

The research was funded by a UKRI Future Leaders Fellowship to DH (MR/S034390/1). We would like to thank Dr. Kai Erdmann, Prof. Sherif ElKhamisy and Prof. Martin Wiedmann for reagents. Microscopy was performed in the Wolfson Light Microscopy Facility using a Nikon wide-field microscope (funded by Medical Research Council SHIMA award MR/K015753/1).This paper was typeset with the bioRxiv word template by @Chrelli: www.github.com/chrelli/bioRxiv-word-template

## Author contributions

DH supervised the study. MEG and DH devised the concept. MEG, MC, AEMI performed experiments. All authors designed experiments and analysed the data. DH and MEG wrote the manuscript.

## Materials and Methods

### Cell culture

Cells were maintained in a humidified incubator (Panasonic, MCO-170AICUV-PE) at 37°C and 5% CO2. Cells were passaged every 2-3 days in their appropriate media. HT1080, NIH/3T3, and IMR90 cells were cultured in Dulbecco’s Modified Eagle’s Medium - high glucose (DMEM; Sigma Aldrich, D6546), and THP1 cells in RPMI1640, HIEC-6 in Opti-MEM Reduced Serum Medium. To make complete growth media for HT1080 and NIH/3T3, and THP1 cells basal media were supplemented with: 2mM L-glutamine, and 10% fetal bovine serum (FBS). For IMR90 cells: 15% FBS, 1% non-essential amino acids (NEAA), and 1% GlutaMAX. For HIEC-6: 20 mM HEPES, 10 mM Glutamax, 10 ng/ml epidermal growth factor and 4% FBS. All base media were supplemented with 10 U/ml Penicillin/Streptomycin (Gibco, 11548876), 50 μg/ml Kanamycin sulphate (BioBasic, KB0286). To differentiate Thp1 cells to macrophages, 100 ng/ml Phorbol 12-myristate 13-acetate PMA was added for 72 hours.

### Recombinant toxin purification and intoxication assays

The typhoid toxin was purified as previously described in Ibler et al 2019. Cells were intoxicated with 20 ng/ml toxin for 2h, washed three times with sterile PBS to remove any extracellular toxin and chased with fresh complete growth media for the duration of the experiment until fixation or conditioned media harvest.

### Treatment with drugs or purified proteins

Cells were treated with 20 μM APH or 10 μM ETP for 24 hours prior to washing 3X with PBS and incubating with fresh complete growth medium. TGF-β RI Kinase Inhibitor III, KU55933 (ATM inhibitor) and AZD6738 (ATR inhibitor) were added at 1 μM after washing the toxin off or in conditioned media. Caffeine was added at 10 mM for 1 hour pre-intoxication, during intoxication, and after removal of the toxin. Purified proteins were added in the following concentrations 10 ng/ml TGFB3, 500 ng/ml Activin A, 200 ng/ml GDF15, 500 ng/ml Wnt5a.

### Knockdown by siRNA transfection

Per well of a 24-well plate format, Lipofectamine RNAiMax (0.25 μl or 0.5 μl) and siRNA (0.5 μl of 20 μM stock) were prepared in two different tubes of 25 μl serum-free, then mixed together for 5 minutes at room temperature with gentle tube flicking. The 50 μl mix of siRNA and lipofectamine were added to 450 μl complete growth DMEM, and incubated on cells for 48h before any further treatments or assays. The final concentration of siRNA in culture was 20 nM.

### *Salmonella* infection assay

*S*.Javiana (S5-0395) and Δ*cdtB* (M8-0540) were kind gifts from Prof. Martin Weidmann (New York). A *Salmonella* colony was cultured at 37 °C in a shaking incubator and used to infect HT1080 cells seeded on glass coverslips. The plate was then spun for 1 min at 1000 × g and incubated for 30 min at 37°C 5% CO2. Cells were then washed with PBS, and incubated with 50 μg/ml gentamicin-containing media (Chem Cruz, sc203334) for 1.5h followed by 10 μg/ml gentamicin for the rest of the experiment. For CFU invasion assays, cells were serum-starved 24h prior to infection to deprive the cells of membrane ruffling stimulants in FBS. Serum-free conditioned media was added for 1h before infection. After 1.5h treatment with gentamicin, cells were lysed with 1% triton X in water for 5-10 mins, and pipetted vigorously. Serial dilution of the lysis buffer was performed and 5 μl spotted on agar plates then incubated overnight in a dry incubator at 37°C. Colonies were then counted manually, and the highest, countable *Salmonella* colonies (i.e. the lowest dilution with visible colonies) were counted and normalised to calculate *Salmonella* colony forming units (CFUs).

### Immunofluorescence

Media was washed off with PBS, then fixed with 4% paraformaldehyde (PFA) in PBS for 10-15 mins at room temperature. Cells were washed two more times with PBS (Biotech, PD8117) then blocked and permeabilised using a 3% BSA (Sigma-Aldrich, 1073508600), 0.2% Triton X-100 (VWR, 28817.295) in PBS, at room temperature for 1h. Primary and secondary antibodies were added in blocking buffer consecutive for 1h and 30min, respectively then washed with PBS then water and left to dry. Coverslips were then mounted and counterstained on 6 μl of VectaShield mounting agent with DAPI (Vector Lab, H1200), and sealed before being imaged on Nikon’s Inverted Ti eclipse equipped with an Andor Zyla sCMOS camera (2560 × 2160; 6.5μm pixels). The objectives used were Plan Apo 10x (NA 0.45); Plan Apo 20x (NA 0.75); Plan Fluor 40x oil (NA 1.3); Apo 60x oil (NA 1.4); Plan Apo 100x Ph oil (Na 1.45); Plan Apo VC 100x oil (NA 1.4). Quad emission filters for used with SpectraX LED excitation (395nm, 470nm, 561nm, 640nm). The imaging software used was NIS elements software.

### Senescence assays

Senescence was assayed by histological staining using Senescence β-Galactosidase Staining Kit (Cell Signalling Technology #9860S), or fluorescence using CellEvent™ Senescence Green Probe (Invitrogen, C10851) as per the manufacturer’s instructions. To assay for cell cycle arrest, Click-iT™ EdU Cell Proliferation Kit for Imaging, Alexa FluorTM 647 dye (Thermofisher, C10340) was used per the manufacturer’s instruction. For example, HT1080 duplication time is 18h, therefore EdU was added to the culture 18-24h before fixation. Where EdU were to be co-stained with CellEvent Green and other immunofluorescence staining, the order of assays was CellEvent Senescence Green, EdU staining Kit, and then immunofluorescence. Histological SA-β-gal was imaged using the inverted microscope Nikon Eclipse Ts2 equipped with sCMOS Ds-Fi-3 camera with built-in Diascopic, high-intensity LED illumination system at 10x and 40x objective lenses.

### Secretome harvest

Conditioned media was centrifuged at 2000-6000 × g for 5 mins to pellet the cells before filtering through 0.2μm filters to prevent cell contamination and release of non-SASP intracellular factors due to the filtration pressure. Conditioned media was stored at −20°C for short term storage and −80°C for long term storage. Before incubation on cells, conditioned media was either supplemented 1:1 with complete growth media, or 10% FBS if conditioned media was serum-free.

### Sample preparation and mass spectrometry

A protease mixture of leupeptin, pepstatin, and chymostatin (Sigma Aldrich, 62070, 77170, C7268, respectively) each at a final concentration of 10 μg/ml) was added to harvested conditioned media immediately. Samples were concentrated using amicon filters with 3 KDa cutoff down to 250 μl at 4000×g using swing bucket centrifuge at 4°C. 50 μl of the concentrated media was used for S-trap digestion. Equal volume of 2x S-trap lysis buffer containing 10% SDS (Sigma-Adrich, 05030), 100mM TEAB, pH 7.55 (Thermofisher, 90114) was added to the samples. Next, the samples were reduced by adding TCEP (Merck, 646547) at a final concentration of 10 mM. The samples were then heated at 70°C for 15 mins at 750 rpm in a shaking incubator then cooled down for 5 mins at room temperature. Next, the samples were alkylated by adding 20 mM iodoacetamide (Merck, I6125) in the dark for 30 mins at room temperature. The samples were then acidified by adding aqueous phosphoric acid to a final concentration of 1.2%. S-trap binding buffer (90% aqueous methanol, 0.1M TEAB, pH 7.1) was added to the solution in a ratio of 1:7. Samples were then loaded into the S-trap columns (Protifi, C02), 150 μl at a time gently and centrifuged at 4000 x g for 10 seconds to pass through the S-trap. The S-trap was then washed 3X with 150 μl binding buffer before transferring it to a clean 2 ml Eppendorf tube. MS grade trypsin (Fisher Scientific, 13464189) resuspended in 0.1% TFA was then added to a ratio of 1:10 trypsin:protein w/w in 50 mM TEAB buffer. 30 μl of trypsin in TEAB buffer was added to the samples and air bubbles were avoided. The S-trap was sealed and incubated at 47°C for 1 hr without shaking. The peptides were then eluted by adding 40 μl of 50 mM TEAB, 40 μl of 0.2% aqueous formic acid (Fisher Chemical, A117-50) then 40 μl of 50% ACN containing 0.2% aqueous formic acid at 4000×g for 10 seconds each. Samples were dried in a SpeedVac (Eppendorf) at 45°C for 60-90 minutes with vacuum setting V-AQ. Peptides were resuspended in 0.5% formic acid and mixed by vortex at the lowest speed for 10 minutes. 18 μl of each sample was analysed by nanoflow LC-MS/MS using an Orbitrap Elite (Thermo Fisher) hybrid mass spectrometer equipped with an easyspray source, coupled to an Ultimate RSLCnano LC System (Dionex). The system was controlled by Xcalibur 3.0.63 (Thermo Fisher) and DCMSLink (Dionex). Peptides were desalted on-line using an Acclaim PepMap 100 C18 nano/capillary BioLC, 100A nanoViper 20 mm × 75 μm I.D. particle size 3 μm (Fisher Scientific) and then separated using a 125-min gradient from 5 to 35% buffer B (0.5% formic acid in 80% acetonitrile) on an EASY-Spray column, 50 cm and N-terminal protein acetylation as variable modifications, cysteine carbamidomethylation as a fixed modification, match between runs enabled with a match time window of 0.7 min and a 20-min alignment time window, label-free quantification enabled with a minimum ratio count of 2, minimum number of neighbours of 3 and an average number of neighbours of 6. A first search precursor tolerance of 20ppm and a main search precursor tolerance of 4.5 ppm was used for FTMS scans and a 0.5 Da tolerance for ITMS scans. A protein FDR of 0.01 and a peptide FDR of 0.01 were used for identification level cut-offs. MaxQuant output was loaded into Perseus version 1.5.6.0 and all LFQ intensities were set as main columns. The matrix was filtered to remove all proteins that were potential contaminants, only identified by site and reverse sequences. LFQ intensities were then transformed using log2(x) default function. To group experimental conditions, rows were categorically annotated with their condition names. Rows with no LFQ intensities in all categorised groups were eliminated by filtering rows based on valid value in at least one group with a minimum number of 3 out of 4 repeats for statistical analysis. Subsequently, data were visualised using multi-scatter plots and Pearson’s correlation analysis and outliers removed. Data were then normalised by subtracting column medians and missing values were randomly imputed with a width of 0.3 and downshift of 1.8 from the standard deviation. In order to identify quantitatively enriched proteins between groups, two-sided Student’s t-tests were performed with a permutation-based FDR calculation (FDR = 0.05) with an (S0) =0.1. Data was then exported into an excel file and input into GraphPad Prism to create the figures and plots presented.

### GeneChip microarray

Cells were seeded in T75 tissue culture flasks for a 30% confluency before intoxication with txWT and txHQ at 5ng/ml as described. After 48 h, cells were trypsinised and RNA isolation was carried out. Samples were analysed at the Sheffield Institute of Translational NeuroscienceI (SITraN (kindly supported by Paul Heath) on Clariom™S assay, human (ThermoFisher Scientific, 902927). Analysis was performed with Transcriptome Analysis Console 4.0 software (Applied Biosystems, Thermo Fisher Scientific).

### Quantification

Manual quantification was carried out using FIJI cell counter plugin. Raw numbers were then recorded in Microsoft Excel. Analysis of the γH2AX and RPA phenotype was carried out using MATLAB image processing code (https://github.com/nbul/Nuclei) (Ibler et al 2019). CellProfiler 4.2.1 was used to semi-automate quantification of proteins. Nuclear components were quantified by either i) nominating an absolute signal threshold before automatic classification within CellProfiler (e.g. EdU-positive, EdU-negative cells), or ii) by measuring intensity and setting a threshold using the upper quartile of the negative control (e.g. 53BP1, γH2AX, pSMAD2). Cytoplasmic components were quantified using global image intensity divided by the number of nuclei counted. SA-β-gal positive cells were identified by expanding an area around the nucleus (around 50 px). To quantify lamin intensity, each nucleus was first thresholded in FIJI and then input into CellProfiler. A ratio between maximum intensity of lamin at the edge of the nucleus to intensity of the centre of the nucleus was calculated and displayed on the chart.

### Image processing

Fiji 2.1.0/1.53c macro code was created to automate processed images. × 50 μm ID, PepMap C18, 2 μm particles, 100 A pore size (Fisher Scientific). Briefly, brightness and contrast are normalised across all images of inter-The Orbitrap Elite was operated with a cycle of one MS (in the Orbitrap) acquired at a resolution of 60,000 at m/z 400, with the top 20 most abundant multiply charged (2+ and higher) ions in a given chromatographic window subjected to MS/MS fragmentation in the linear ion trap. An FTMS target value of 1e6 and an ion trap MSn target value of 1e4 were used with the lock mass (445.120025) enabled. Maximum FTMS scan accumulation time of 500 ms and maximum ion trap MSn scan accumulation time of 100 ms were used. Dynamic exclusion was enabled with a repeat duration of 45 s with an exclusion list of 500 and an exclusion duration of 30 s.

### LC-MS/MS analysis

All raw mass spectrometry data were analysed with MaxQuant version 1.6.10.43. Data were cross-referenced against a human UniProt sequence database (June 2015) using the following search parameters: digestion set to Trypsin/P with a maximum of 2 missed cleavages, methionine oxidation est, a pre-set ROI is used to crop images to regions of interest, and a pre-set scale bar is added. DAPI nuclei are then outlined, overlaid with the other channels and saved as png files. Many variants of the code have been created to cater for 2-channel, 3-channel and 4-channel images, with either normal composite images or with DAPI outlines. The codes can be found via this link: https://drive.google.com/drive/folders/1SwUO-bgyW0rjY9MetAfKGtdd77FsCahF?usp=sharing

### Statistics

Data was recorded and organised in Microsoft Excel. Graphs were generated and statistics were performed using GraphPad Prism 9.0.0. Statistical significance was denoted using asterisks (*) where *p<0.05, **p<0.01, ***p<0.001, ****p<0.0001, ns denotes non-significance. Error bars indicate standard error of the mean (SEM). Adobe Illustrator was used to manually hand-draw illustrations and assemble all results figures.

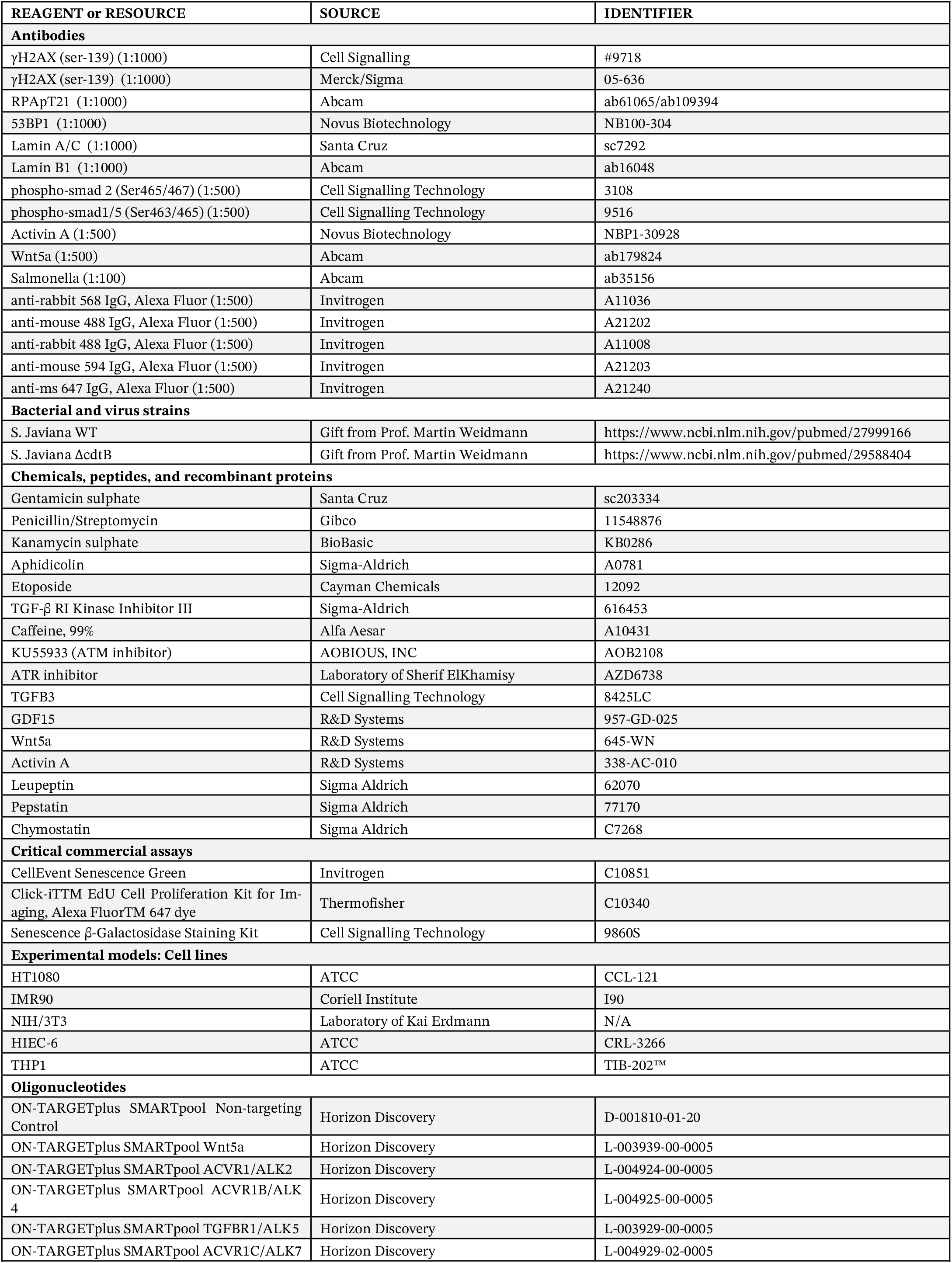
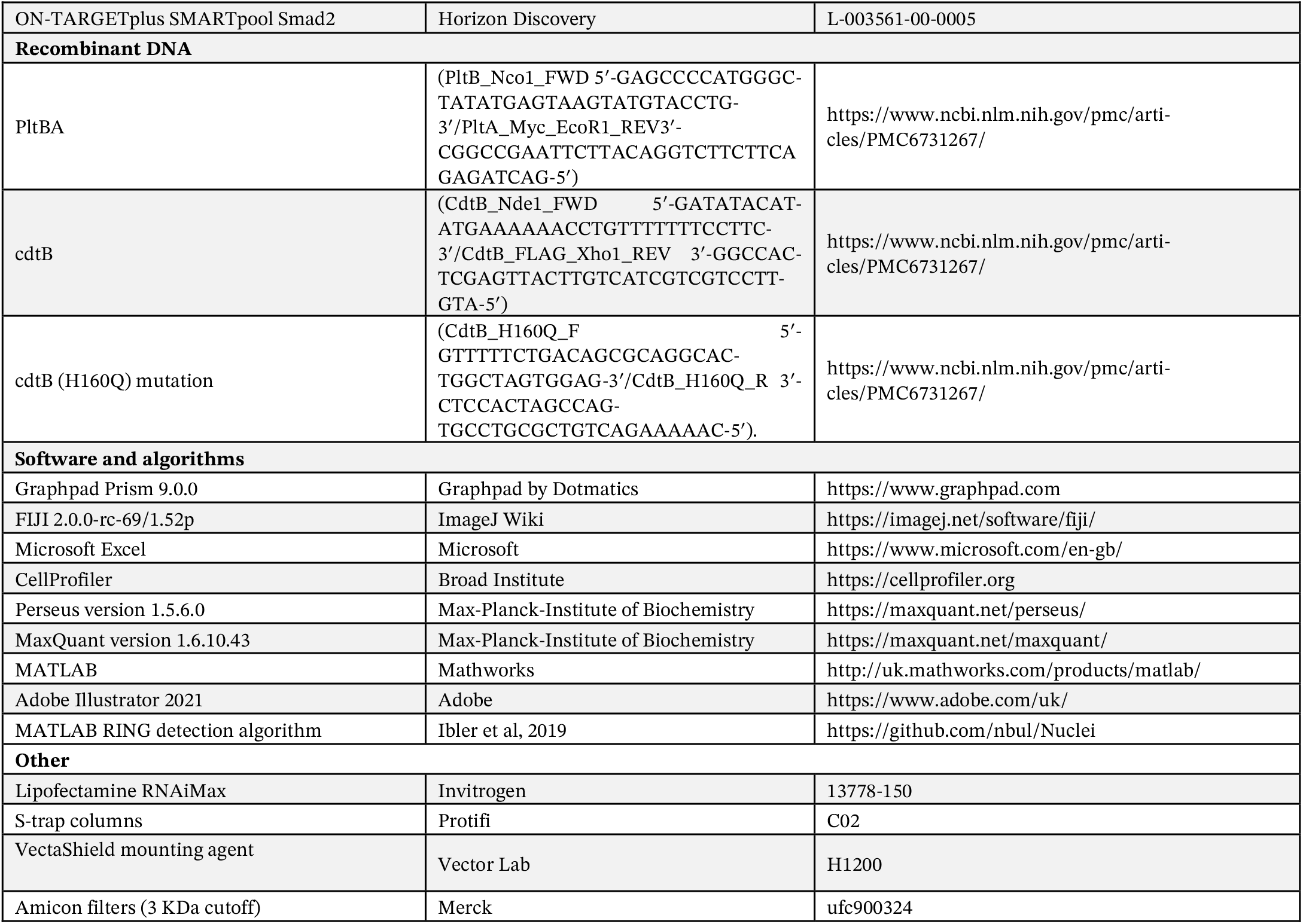

**Figure S1.**
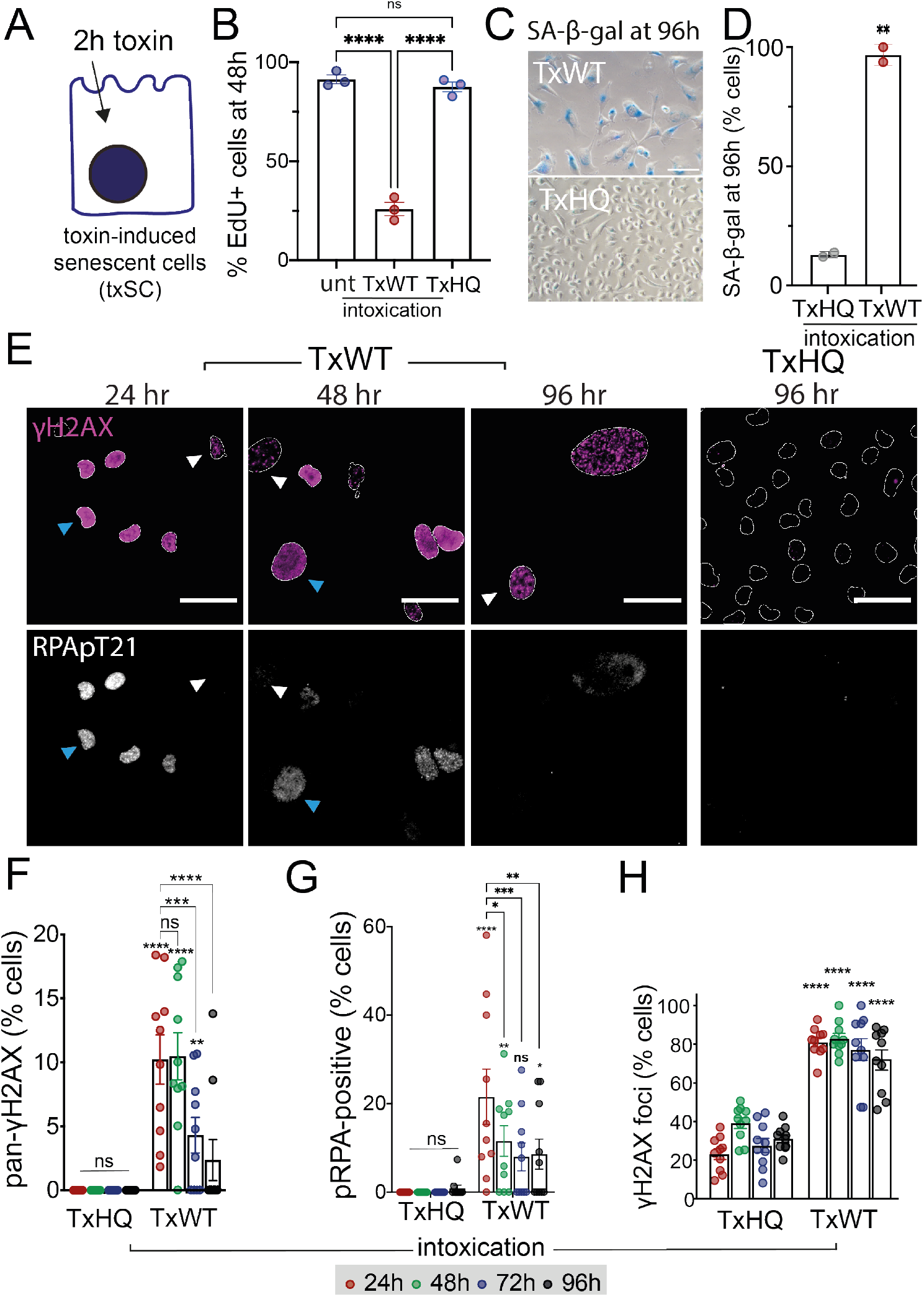
Evolution of DDRs in cells undergoing toxin-induced senescence. (**A**) Cartoon depicting toxin-induced senescent cells (TxSC). (**B**) Percentage of EdU-positive cells at 48h in untreated cells (unt) or cells treated for 2h with TxWT or TxHQ. (**C**) Representative images of SA-β-gal at 96h in cells treated as (B). (**D**) Quantification of the experiment in (C) where circles indicate 2 biological replicates. (**E**) Representative images of γH2AX and replication stress marker RPA phosphorylated at T21 (RPApT21) at indicated time-points in cells treated as (B). Outlines of DAPI-stained nuclei shown. Scale bars 50 μm. Quantification of phenotypes shown in (E): (**F**) Percentage of nuclei with pan-γH2AX, (**G**) Percentage of nuclei with phosphorylated RPA, and (**H**) Percentage of nuclei with γH2AX foci. Circles indicate technical replicates from 2 biological replicates. Asterisks indicate significance (*p<0.05, **p<0.01, ***p<0.001, ****p<0.0001, ns=no significance) calculated by (B) One-way ANOVA test with Tukey’s multiple comparison, (D) unpaired t-test, (F-G) Two-way ANOVA with Tukey’s multiple comparison test, and (H-I) Two-way with Sidak multiple comparisons tests. Error bars SEM. **Fig S1** connected to **Fig 1**.

**Figure S2.**
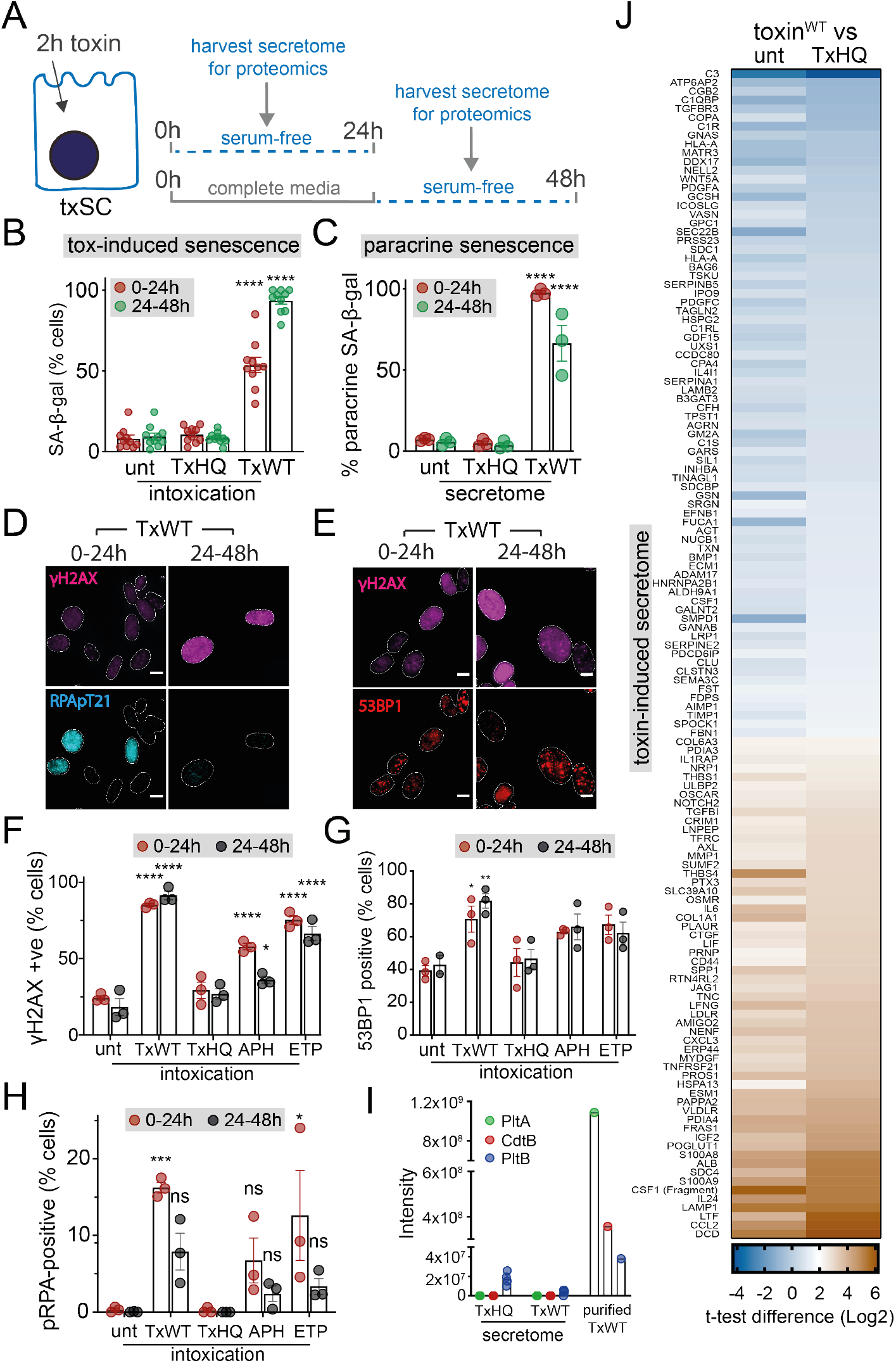
Toxin-induced responses in serum-free conditions. (**A**) Schematic depicting experimental approach for harvesting secretome from TxSCs prior to proteomics. (**B**) Quantification of toxin-induced senescence at 24h or 48h. SA-β-gal in TxSCs depicted in (A) following no treatment (unt) or 2h treatment with TxWT or TxHQ assayed at 24h (i.e. endpoint for harvesting secretomes accumulating between 0-24h) or 48h (i.e. endpoint for harvesting secretomes accumulating between 24-48h). Circles represent technical replicates from 1 biological replicate. (**C**) Quantification of paracrine senescence at 96h. Secretomes were harvested, as depicted in (A), from untreated (unt), TxWT- or TxHQ-treated cells shown in (B) before supplementation with 10% FBS then incubation with naive cells and assaying SA-β-gal at 96h. Circles represent 3 biological replicates. (**D**) Representative images of γH2AX (magenta) and RPApT21 (cyan) from the same experiment as (B). (**E**) Representative images of γH2AX (magenta) and 53BP1 (red) from the same experiment as (B). Quantification of phenotypes at 24h or 48h from the experiment in (B) with additional treatments APH and ETP: (**F**) Percentage of γH2AX-positive cells, (**G**) Percentage of 53BP1-positive cells, and (**H**) Percentage of pRPA-positive cells. Circles represent 3 biological replicates. (**I**) LC-MS/MS peptide intensity identified PltB in secretome^TxWT^ (TxWT) and secretome^TxHQ^ (TxHQ). Purified TxWT used as positive control. Circles indicate 4 biological replicates. (**J**) Heat map listing TxWT-induced host cell secretome relative to secretome^unt^ (unt) and secretome^TxHQ^ (TxHQ). Sorted from highest to lowest t-test difference relative to secretome^TxHQ^. Blue bars indicate protein up-regulated in secretome^TxWT^ and brown indicates down-regulated. Asterisks indicate significance (*p<0.05, **p<0.01, ***p<0.001, ****p<0.0001, ns=no significance) calculated by (B-H) Two-way ANOVA with (B-C) Dunnetts and (F-H) Sidak multiple comparisons test. Error bars SEM. Scale bars 50 μm in (D-E). **Fig S2** connected to **Fig 1**.

**Fig S3.**
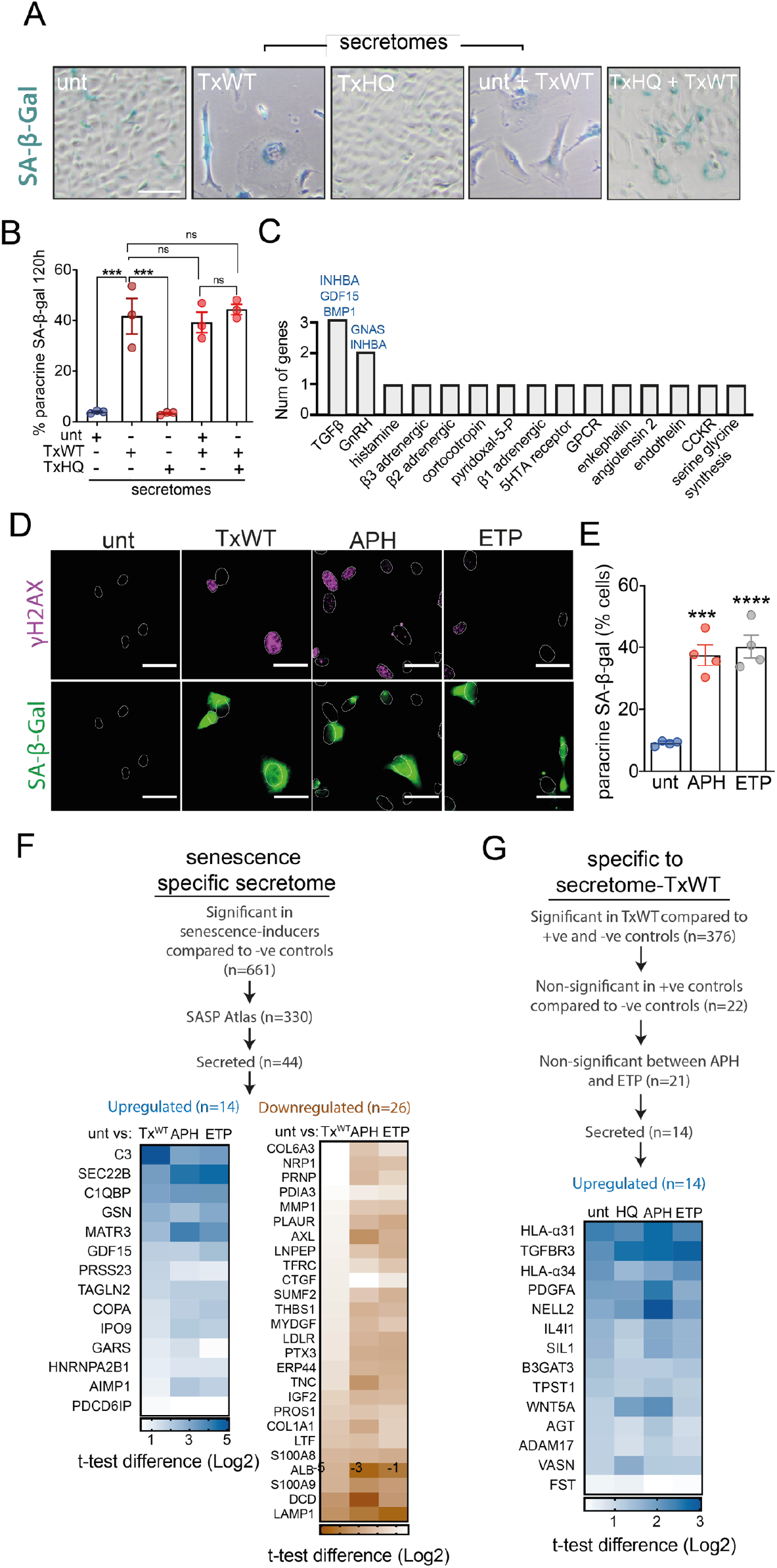
Comparative analysis of SASP induced by typhoid toxin, aphidicolin and etoposide. (**A**) Paracrine senescence mediated by upregulated proteins in secretome^TxWT^. Naive cells treated with indicated combinations of secretomes supplemented with 10% FBS before assaying for SA-β-gal at 120h. (**B**) Quantification of the experiment shown in (A). Circles represent 3 biological replicates. (**C**) Panther pathway analysis of proteins in the secretome^TxWT^ listed in Fig 1H. Blue text indicates up-regulated genes. (**D**) Representative images of γH2AX and SA-β-gal in untreated cells or cells treated with TxWT, APH or ETP at 96h. Outlines of DAPI-stained nuclei shown. Scale bars 50 μm. (**E**) Paracrine senescence by secretome^APH^ and secretome^ETP^. Cells treated for 24h with APH or ETP were incubated for a further 48h in the absence of drugs before harvesting secretome^APH^ and secretome^ETP^. Naive cells were treated with secretome^APH^ or secretome^ETP^ before assaying SA-β-gal at 96h. Circles represent 4 biological replicates. (**F**) Identified proteins common amongst the secretomes of TxWT-, APH- and ETP-treated cells relative to untreated. (**G**) Identified proteins specifically enriched in secretome^TxWT^. Flowcharts in (F) and (G) outline steps used to filter significant proteins. Asterisks indicate significance (***p<0.001, ****p<0.0001, ns=no significance) calculated by (B, E) One-way Anova with Tukey’s multiple comparison test. Error bars SEM. **Fig S3** connected to **Fig 1** and **Fig 2**.

**Figure S4.**
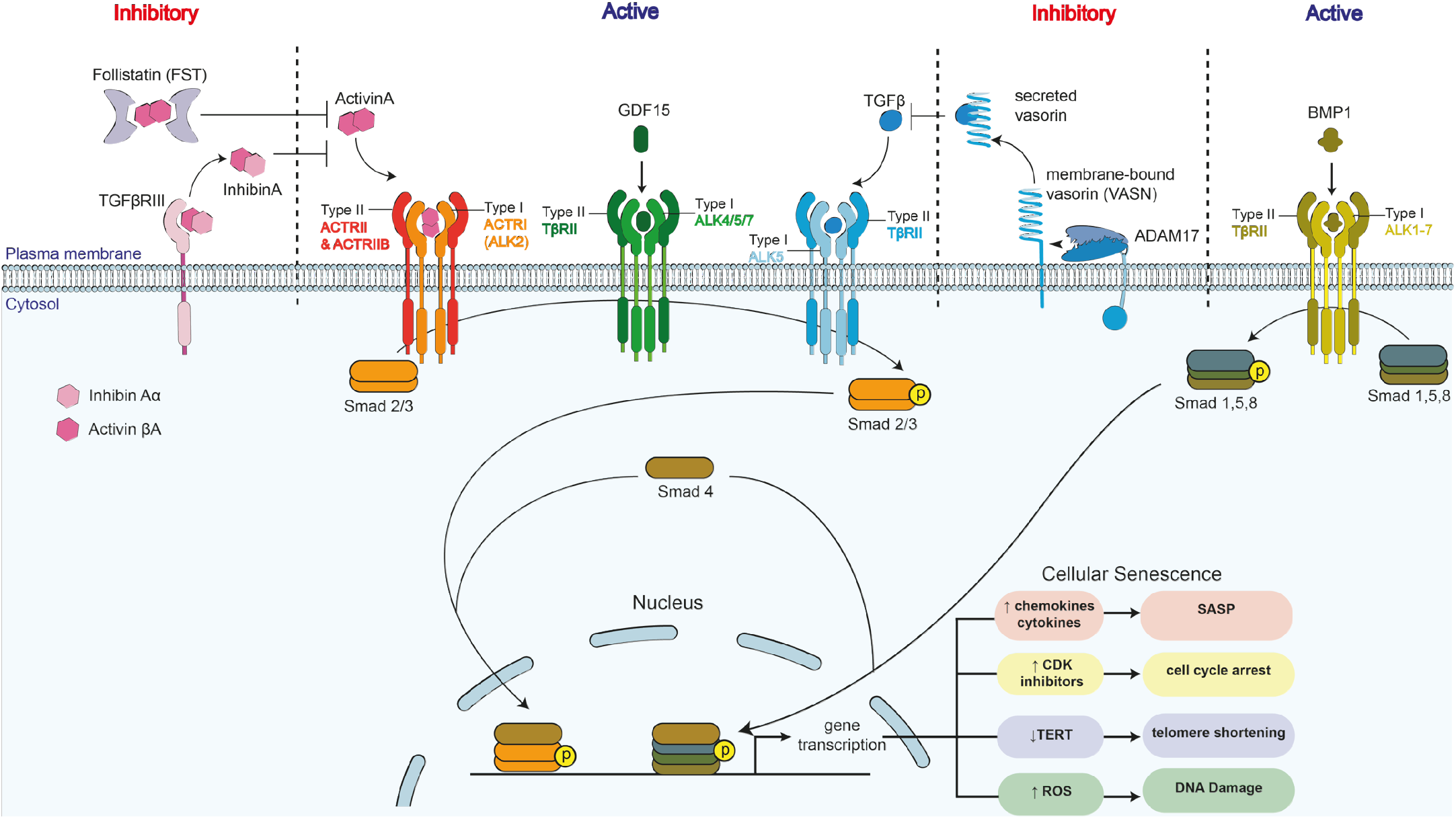
Cartoon summarising TGFβ pathway. Schematic representation of secretome^TxWT^ proteins implicated in the TGFβ pathway. The different players identified are either involved in activating or inhibiting TGFβ signalling. Activation: ligands Activin A, TGFβ, GDF15 or BMP1 bind to their respective Type I/II tyrosine kinase receptors causing phosphorylation and activation of SMAD transcription factors(36). Phosphorylated SMADs translocate to the nucleus to drive transcription of genes involved in senescence. Inhibitory: follistatin and TGFBRIII sequester Activin A while the metalloprotease ADAM17 liberates the ectodomain of cell surface protein vasorin, which sequesters TGFβ. These activities reduce SMAD activation.

**Figure S5.**
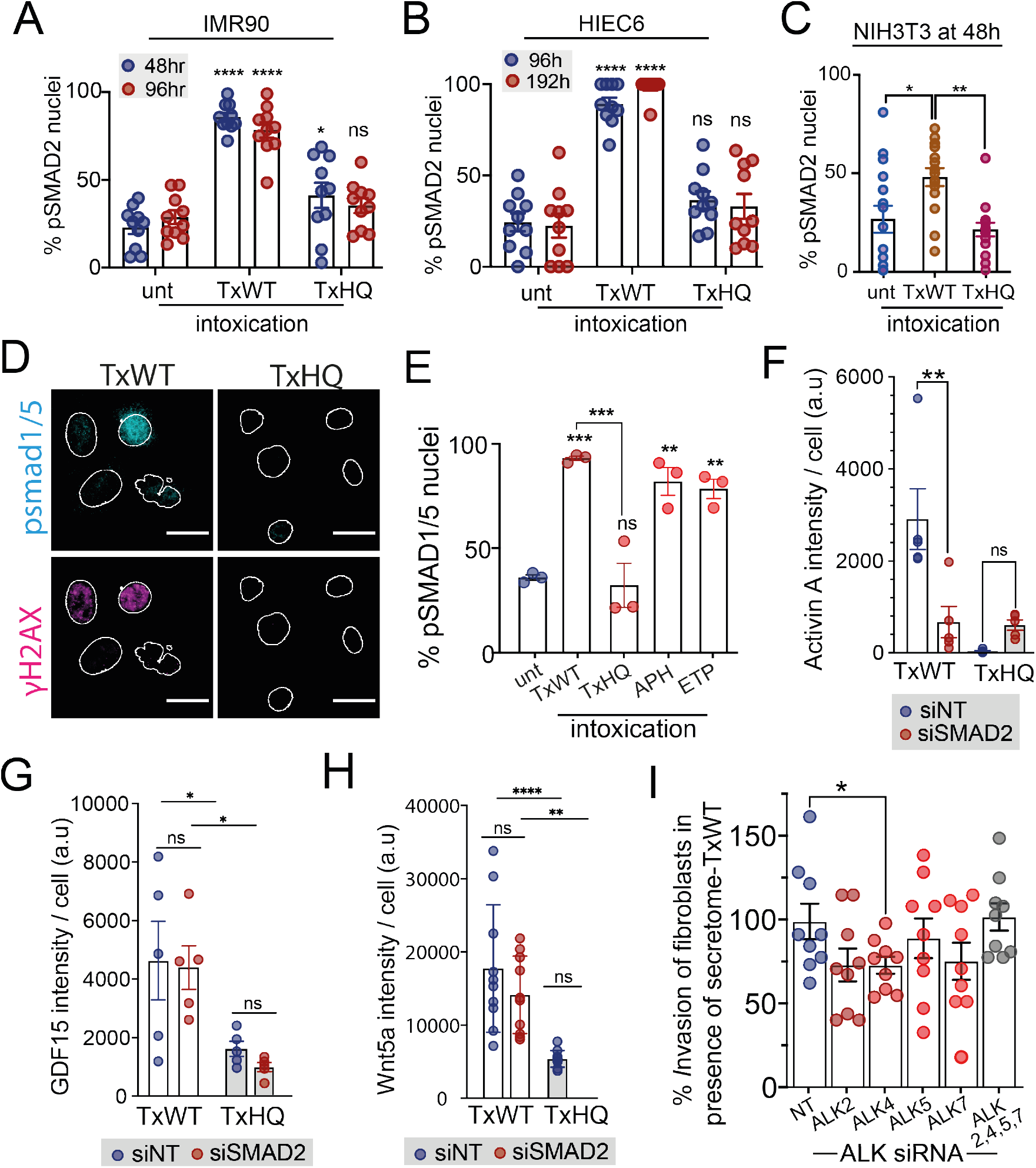
TGFβ signalling in cells treated with typhoid toxin, aphidicolin and etoposide. Percentage of pSMAD2-positive nuclei at indicated time points following no treatment (unt) or treatment with TxWT or TxHQ in the following cell lines: (**A**) IMR90 fibroblasts, (**B**) HIEC6 epithelial cells, and (**C**) NIH3T3 fibroblasts. Circles represent technical replicates (1 biological replicate). (**D**) Representative images of pSMAD1/5 (cyan) and γH2AX (magenta) in TxWT- or TxHQ-treated HT1080 cells at 48h. Outlines of DAPI-stained nuclei indicated. Scale bar 25μm. (**E**) Quantification of (D) with additional treatments APH and ETP. Circles represent 3 biological replicates. Quantification of (**F**) Activin A expression, (**G**) GDF15 expression, or (**H**) Wnt5a expression at 48h in siNT- or siSMAD2-transfected cells treated with TxWT or TxHQ. Circles represent technical replicates (1 biological replicate). (**I**) *Salmonella* invasion into non-phagocytic cells. IMR90 fibroblasts were transfected with siRNAs targeting indicated ALK receptors before 30 min infection with *S*.Javiana in the presence of secretome^TxWT^. Relative invasion was calculated by culturing *Salmonella* CFUs from infected cells. Circles represent technical replicates (3 biological replicates). Asterisks indicate significance (*p<0.05, **p<0.01, ***p<0.001, ****p<0.0001, ns=no significance) calculated by (A, B) Two-way ANOVA with (A) Tukey’s multiple comparison, or (B) Dunnett’s multiple comparison, (C, E) one-way ANOVA with Dunnett’s multiple comparisons, (F, G, H) two-way ANOVA with Sidak multiple comparisons test, and (I) unpaired t-test. Error bars SEM. **Fig S5** connected to **Fig 3** and **Fig 4**.

